# SynLS: A novel diffusion-transformer framework for generating high-quality wearable sensor time series data to enhance health monitoring

**DOI:** 10.1101/2025.05.11.653212

**Authors:** D. Lin, Y. F. Ji, J. A. A. McArt, J. Li

## Abstract

While global medical research is poised to benefit from the rapid advance of artificial intelligence (AI) technologies, veterinary medicine research often faces significant limitations due to data scarcity and availability issues. To address this issue, we proposed a generative modeling framework, SynLS, for generating highly realistic synthetic wearable sensor data. Leveraging diffusion architecture and transformer encoder mechanism, SynLS addressed the intricate challenges posed by these real-world wearable sensor data, including varied length, multiple dimensions, high diversity, high noise, periodicity, and trend. We have validated SynLS on four publicly-available livestock wearables databases with records for three health events (calving, estrus and diseases), and demonstrated its ablility in producing high-fidelity wearable sensor data, which could improve the downstream health events prediction tasks by 18.5% and 26.8% under two evaluation scenarios based on instance and timestamp, respectively. Additionally, introducting raw tri-axial accelerometer databases collected from animals and human further demonstrated extensibility of our framework, significantly enhancing downstream behavior classification tasks by 38.8% and 83.8%, respectively. The technical framework proposed in this work offers a potential generalized solution for data supplementation in wearables sensor databases, with potential applicability across veterinary medicine and other medical domains facing resource constraints.

## 1. Introduction

Ground-breaking advancements in intelligent wearables have not only revolutionized human health monitoring strategy, but also reshaped livestock health monitoring programs in the non-invasive and real-time manner ^1,2,3^. Most of these wearables transform accelerations into behavioral and physiological parameters, enabling precise description of feeding, activity, and rumen conditions ^4^. Although the changing patterns of these continuous parameters show potential for alerting producers to heat events and health disorders, they come with limitations of high false positive and negative rates ^4,5^. These challenges could be addressed by the secondary uses of the vast amounts of continuous data and a wide range of machine learning (ML) models. Yet, in practice, these wearables data are rarely shared beyond the commercial organization that initially collected the data, which hamper efforts for model development, clinical research and practical applications. To accelerate Artificial Intelligence (AI) applications in veterinary medicine, it’s necessary to develop methods to synthesize ‘realistic’ time series data from these intelligent wearables.

Synthetic high-quality time series data should typically present two key characteristics: 1) high fidelity, meaning the synthetic data must closely resemble real sensor data collected from animals; and 2) high utility, indicating that the synthetic data should be helpful for downstream tasks such as predicting important health events. Achieving these characteristics presents several challenges. First, different wearable devices generate varying resolutions of time series, with high-resolution units capturing detailed patterns like circadian rhythms ^6,7^ and necessitating longer sequences for comprehensive record of information^8^. Second, wearable devices typically produce multi-dimensional data (e.g., multiple behavioral/ physiological time series indices) ^9^, making it essential to capture their interdependencies during the generation process. Additionally, the animal behaviors are highly diverse, influenced by breed, age, parity, environment, season, and management practices ^10,11,12^, leading to complex patterns underlying wearable sensor data across individual animal and farms. Finally, these sensor-recorded parameters could serve as early indicators of significant health events ^13,14,15^, requiring the learning of underlying patterns and trends from the past information.

The generation of synthetic real-world medical time series data has received increasing attention with a predominant focus on generating high-dimensional and multi-faceted data. Some conventional generative methods, such as anomalies injection ^16^ and Dynamic Time Warping Barycentric Averaging (DBA) ^17^, struggle to capture multidimensional distribution across variables and temporal dynamics across time, leading to their gradual obsolescence. In recent years, generative AI offers new horizons, particularly for synthesizing continuous electronic health records (EHR) ^18,19,20^ and neurophysiological data, such as neural population dynamics ^22^, neural spike trains ^23^, and electroencephalography (EEG) ^24,25,26^, by utilizing the power of Generative Adversarial Networks (GANs) and Variational AutoEncoders (VAEs). Despite this, the main drawbacks of current models are the commonly known pitfalls of GANs and VAEs, including model collapses due to unstable training regimes ^27,28^, and blurry generation because of injected noise and imperfect reconstruction ^29^. Furthermore, although GANs-based ^30^ and VAEs-based ^31^ models have been presented for the synthetic generation of real-world time series, the varied length, multiple dimensions, high diversity, high noise, and periodicity and trend of wearable sensor data is usually not taken into account for the generation. Specifically, to the best of our knowledge, time series synthesis has not been yet explored for wearables in veterinary medicine field.

In this study, we introduced Denoising Diffusion Probabilistic Models (DDPMs, or diffusion models for short)^32^ into wearables time series generation. Diffusion models offer enhanced flexibility with the form of the target distribution in capturing dynamics and understanding the broader context of signal structures ^33^. The objectives of this study were to (1) generate realistic time series data from on-farm animals wearable sensors using generative diffusion models, to (2) assess whether the synthetic data exhibit high fidelity and utility, and to (3) investigate the extensibility of the model to raw tri-axial accelerometer data from both animals and humans. The motivation of this study was to lower the barriers and costs for the acquisition of wearable sensor time series data, especially collected from on-farm animals, thereby opening new possibilities for health monitoring research and adding novel insights into the power of AI for veterinary medicine and even human medicine.

## 2. Materials and Methods

### 2.1 Dataset description

We included a total of four publicly accessible livestock wearables databases and four publicly accessible tri-axial accelerometer databases (two for livestock and two for human) as extension validation in this study. The locations and time series for these databases are shown in Figure 1.

**Figure 1.**
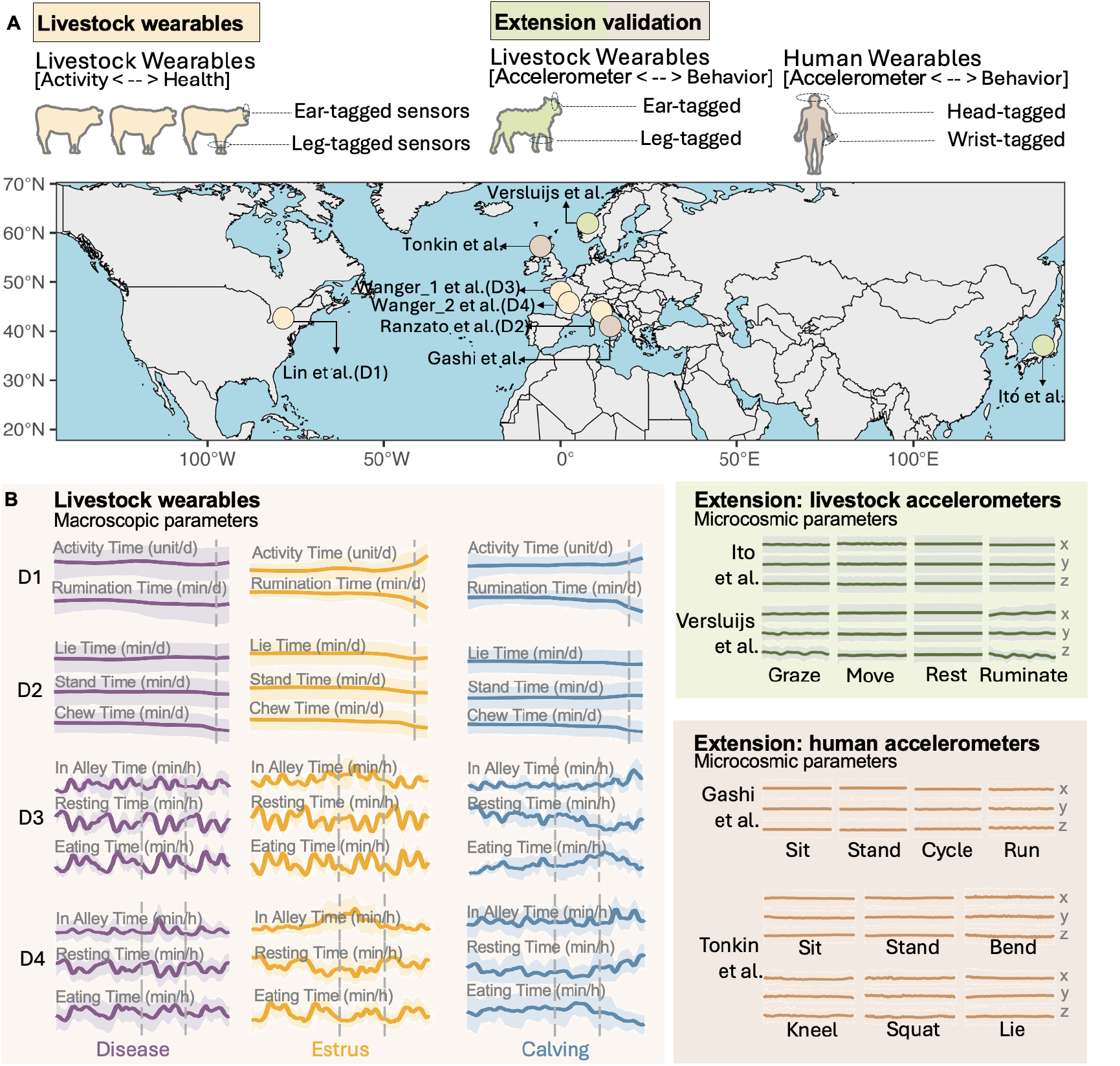
Sample information. A. The locations where eight public wearable sensors databases, including four livestock wearables databases, and four extended tri-axial accelerometer databases, were collected. B. Time series (mean ± standard deviation) for each health event in four livestock wearables databases, and each behavior in four extended tri-axial accelerometer databases. The mean value of the TS is plotted by the solid line, with the shaded area indicating ±1 standard deviation.

#### Livestock wearables databases

The following four databases with defined health events (estrus, calving and diseases for cows) were used for describing the characteristic of on-farm wearable sensor time series data and evaluating both fidelity and utility of our proposed diffusion generative model:

- **Lin et al. (D1)** ^34^ — a low-resolution database comprised of daily activity and rumination data recorded by a neck-mounted or ear-mounted electronic activity and rumination monitoring tag (both SCR Dairy, Netanya, Israel). This database was collected from 185 Holstein-Friesian cows at a commercial dairy farm in Cayuga County, New York, USA, from March 2021 to March 2022.
- **Ranzato et al. (D2)** ^35^ — a low-resolution database consisting of behavioral data regarding daily lying, chewing, and activity times recorded by an ear-tag-based accelerometer (Smartbow GmbH, Weibern, Austria). This database was collected from 369 Holstein-Friesian cows at a dairy farm located in the Po Valley, Italy, from August 2019 to September 2021.
- **Wanger_1 et al. (D3)** ^36^ — a high-resolution database containing behavioral data including hourly eating, resting, and in-alleys times. This database was recorded by CowView system neck collars (GEA Farm Technology, Bonen, Germany) attached to 28 Holstein-Friesian cows in the INRAE Herbipôle experimental farm in France from October 2018 to April 2019.
- **Wanger_2 et al. (D4)** ^36^ — The database presents the same format as the D3 database, but it was collected from 300 Holstein-Friesian cows in the INRAE Herbipôle experimental farm in France from December 2014 to December 2015.

The wearable sensors time series underwent pre-processing to be deployed in the generative model. Following Wagner et al. ^6^’s suggestion, each type of time series was first extracted based on the recorded label from farms, selecting a specific number of days prior to and one day following each health event associated with modified livestock behavior. We excluded subsequent days after the identified health event from our analysis for two reasons: 1) insufficient information to determine whether the behavior was altered; 2) our major focus for the utility was on the early/timely prediction of health events. Min-Max normalization approach was then applied in these sensors data to adjust each dataset’s values within the range of [0, 1]. Only the time series without any missing value was retained for following analysis. The specific information of these four public livestock wearables databases was present in Supplemental Table 1 and Supplemental Figure 1. The detailed information in pre-processing was described in Supplementary Methods 1.

We tested if the time series from four public livestock wearables datasets exhibited 1) non-stationarity using both Box-Ljung test (p-values below 0.05 mean that we can reject this hypothesis at 5% significance and provide evidence of a trend); 2) autocorrelation for dataset D1 and D2 using autocorrelation function (ACF) with the lag between 0 and sequence length n-1 (bars extend beyond 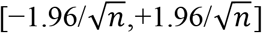 indicate the autocorrelations at these lags are significantly difference from zero); 3) tend and seasonality (per 24h) for datasets D3 and D4 using STL (Seasonal and Trend Decomposition Using LOESS) method ^37^; 4) and multivariate correlations using Spearman’s correlation analysis. The pre-processed time series as well as their characteristics in each database were present in Supplemental Figure 2.

#### Tri-axial accelerometer databases

The following four databases with defined livestock or human behaviors were used to evaluate the utility of our proposed diffusion generative model in raw wearables time series (high resolution (> 1Hz) accelerometry data) which is challenging owing to intricate behaviors, extraneous activity disturbances and substantial individual variability ^38,39,40^. The specific information of these four public tri-axial accelerometer databases was present in Supplemental Table 2 and Supplemental Table 3.

- **Ito et al**.^41^ — a tri-axial accelerometer database with corresponding labeled cow behaviors (grazing, moving, resting and ruminating). This database was obtained with a 16bit +/-2g Kionix KX122-1037 accelerometer (25Hz) attached to the neck of 6 different Japanese Black Beef Cows at a cow farm of Shinshu University in Nagano, Japan in June 2020. We selected the window size of 2.56 seconds, as suggested by the authors ^42^, to get the non-overlapping tri-axial time series with the length of 64 points for modeling.
- **Versluijs et al**.^43^ — a tri-axial accelerometer database relating to different labeled cow behaviors (grazing, moving, resting and ruminating). This database was gained with the Nofence virtual fence collar (10Hz) attached to the neck of 38 Beef Cows (Hereford, Limousine, Charolais, Simmental/NRF/Hereford mix) free-ranging in forested areas in the Innlandet county of Norway during May and June, 2021. We selected the window size of 5.00 seconds to get the non-overlapping tri-axial time series with the length of 50 points for modeling.
- **Gashi et al**. ^44^ — a tri-axial accelerometer database, as a part of Wearable Human Energy Expenditure Estimation (WEEE) ^45^, collected using Muse S headband (52Hz) placed on heads from 17 participants under different physical activities (siting, standing, cycling, and running) in Italy on 2021. We selected the window size of 3.00 seconds to get the non-overlapping tri-axial time series with the length of 156 points for modeling.
- **Tonkin et al**. ^46^— a tri-axial accelerometer database recorded with the wrist-worn sensors (20Hz) from 10 participants under different physical activities (siting, standing, benting, kneeling, squating, lying). This database, as the subset of the dataset for the EPSRC-funded Sensor Platform for HEalthcare in Residential Environment (SPHERE) Challenge ^47^, was collected in the ‘SPHERE House’ in Bristol, UK, during 2016. We selected the window size of 3.00 seconds to get the non-overlapping tri-axial time series with the length of 60 points for modeling.

### 2.2 Architecture of the proposed model SynLS

We developed a transformer design within the diffusion generative framework, termed SynLS. SynLS can be factorized into two key components: (1) a diffusion generative framework for generating synthetic livestock wearable time series; (2) a transformer-based denoising network comprising an encoder with self-attention and positional encoding for capturing the spatiotemporal dynamics and a decoder for reconstruction. The illustration of the framework and model is shown in Figure 2A.

**Figure 2.**
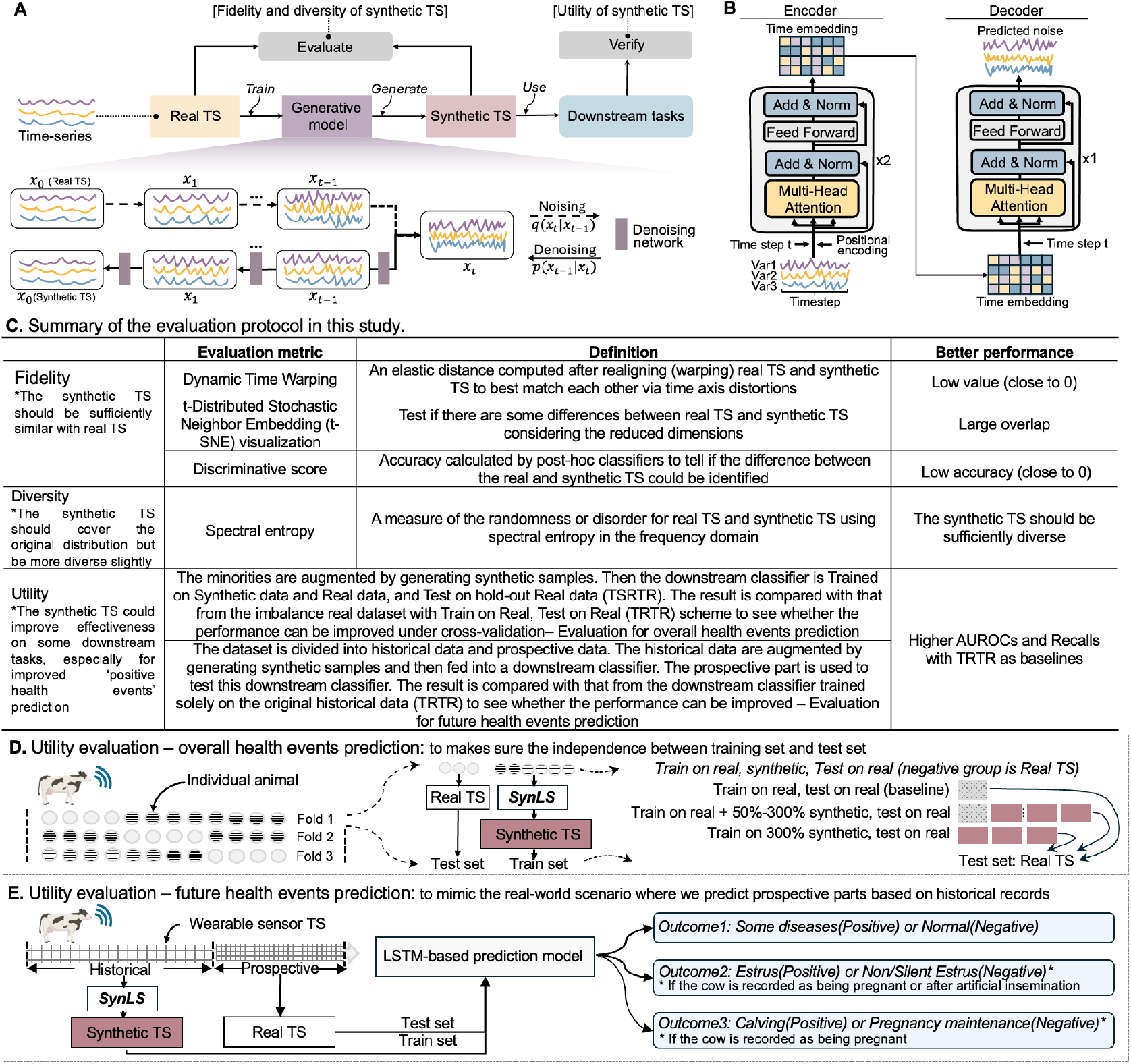
Overall schematics. A. Overall workflow: train a generative model on real time-series data; generate synthetic time-series data; evaluate the synthetic data; finally, verify the effectiveness of synthetic data in some downstream tasks. In this study, DDPM was used as the framework for our generative model, SynLS. B. Illustration of our transformer-based denoising network within SynLS, comprising an encoder with self-attention and positional encoding for capturing the spatiotemporal dynamics and a decoder for reconstruction. C. Summary of the evaluation protocol (fidelity, diversity and utility) used in this study. D. Experimental setup for the first utility evaluation scenario – overall health events prediction to makes sure the independence between training set and test set. E. Experimental setup for the second utility evaluation scenario – future health events prediction to mimic the real-world scenario where we predict prospective parts based on historical records.

#### Diffusion framework for livestock wearable time series generation

Diffusion models are a family of generative models that include a “forward diffusion process”, which progressively converts a data point *x*_0_ into noise, and a “reverse denoising process”, which reverses this approximate diffusion process and therefore generates new data. In this study, we built the diffusion framework based on DDPM ^32^. We concisely summarized the framework here, and more details, including the brief introduction of diffusion models, the related work and the Formulation of DDPM, were provided in Supplementary Methods 2.

In the forward process, we considered a Gaussian diffusion: the conditional distribution of the intermediate state for a time series *x*_*t*_ given the previous state *x*_*t*−1_ is defined by the multivariate normal distribution, if at a time step *t* = 0, 1, 2, · · · *T*:

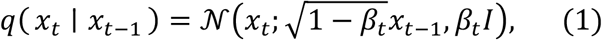

where *I* represents the identity matrix having the same dimensions as the input/original time series *x*_0_, parameter *β*_*t*_ *ϵ* [0,1) represents the variance schedule of the noise level across diffusion steps, and 𝒩(*x*; *μ, σ*) represents the Gaussian distribution of *x* with the mean of *μ* and the covariance of *σ*, respectively. The noise model is Markovian, which allows the distribution of the intermediate state for a time series *x*_*t*_ given *x*_0_ to be expressed as follows:

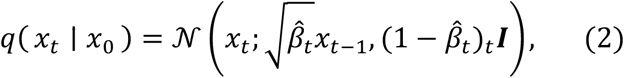

where 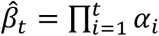 and *α*_*t*_ = 1 − *β*_*t*_. This expression shows that we could produce any intermediate noisy time series *x*_*t*_ via a single step by using the input/original time series *x*_0_ and the variance schedule *β*_*t*_· That is, *x*_*t*_ could be sampled from *q*(*x*_*t*_ ∣ *x*_0_) as follows:

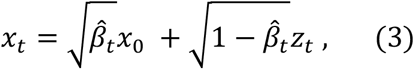

where *z*_*t*_ ∼ 𝒩(0, ***I***)·

The reversed denoising process implements another Markov chain, which transforms the noisy time series *x*_*t*_ back to its original state *x*_0_ from *q*(*x*_0_) via a prior distribution *ρ*(*x*_*t*_) = 𝒩(*x*_*t*_; 0, ***I***) and a learnable transition kernel 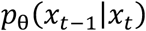.The generative transition distribution is described as follows:

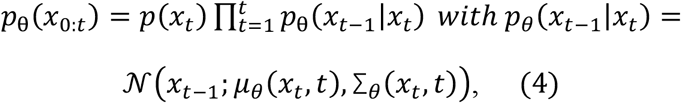

where *θ* denotes model parameters, and the mean *μ* _*θ*_ and variance Σ_*θ*_ are parameterized by neural networks. To train the neural networks such that *p* _*θ*_ (*x*_0_) learns the true data distribution *q*(*x*_0_), the variational lower bound on negative log-likelihood should be optimized, resulting in a simplified training objective:

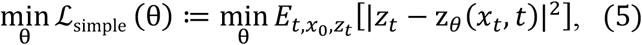

where *E* is the expected value, and z _*θ*_ (*x*_*t*_, *t*) is the network predicted noise from time series *x*_*t*_ at time step t.

Finally, when the neural network is trained, new time series *x*_*new*_ could be generated by first sampling a noisy version *x*_*t*_, then iteratively eliminating noises in *ρ*(*x*_*t*_) = 𝒩(*x*_*t*_; 0, ***I***) from the learnable transition kernel 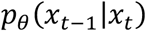,and eventually sample 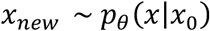.

#### Transformers for feature extraction and reconstruction

The denoising networks consist of two modules based on transformers: an encoder and a decoder.

Encoder. The transformer has two repeated encoder layers, which takes the noisy version *x*_*t*_ as input and converts it into a time embedding matrix *x*_*time*_ that consists of fixed-sized latent embedding vectors representing the temporary dependency of *x*_*t*_. Before *x*_*t*_ is fed into the encoder module, it is first linearly projected into the dimension of *d* = 16, which becomes the value embedding. As transformers can’t interpret the order of time series by default, a fully learnable position embedding with the dimension of *d* = 16 is added to the value embedding. Furthermore, denoising time step *t* is specified using the sinusoidal position embedding, which is defined as below:

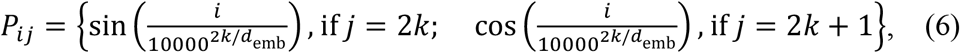

By adding the time step embedding, the input embedding for time series is information-enriched, which contains the embedding of time value, the position of each time point, as well as the denoising time step. This embedding could be recognized in the following self-attention layers. A single self-attention head consists of three learnable matrices: query matrix, a key matrix, and a value matrix. Each matrix shows the dimensionality with *d*_*model*_ × *d*_*head*_, where *d*_*model*_ is the dimensionality of embedding layers, and *d*_*head*_ is the dimensionality of the self-attention head. Input time embedding is multiplied by each of these three matrices independently, producing a query *Q*_*time*_, key K_*time*_, and value vector V_*time*_ for each time point. To determine the contextualized representation for a given time point, the self-attention operation takes the dot product of that query vector with all key vectors. The resulting values are then scaled and softmaxed, producing the “attention weights” for that time point. The principle of the self-attention module is described as follows:

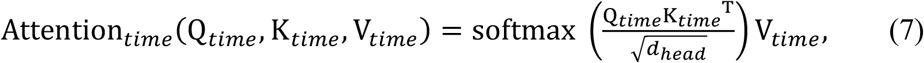

The self-attention operation is independently executed by multiple heads (=2). The output vectors from each head, each having the same dimension, are then concatenated in parallel, which allows their meanings to contextualize each other. Subsequently, the input embedding for time series is added to the multi-head self-attention output via residual connection, and then they are layer-normalized. The result is then passed through a linear projection layer to a dimension of 64, followed by a final linear projection to the dimension of 16. After another residual connection and layer normalization, the time embedding matrix *x*_*time*_ is finally produced.

Decoder. The embedding described above contains the latent representation derived from the noisy version of time series *x*_*t*_ at time step t. The key function of the reconstruction decoder is to recover the noise that is added to *x*_*t*_ from this latent representation. In this step, the denoising time step embedding is concatenated to the latent representation to achieve the retention of time information.

### 2.3 Evaluation of synthetic time series

To comprehensively assess the quality and usefulness of the synthetic livestock wearable sensor time series, we utilized a systematic evaluation framework with respect to three key metrics: 1) fidelity, which measures how closely the synthetic time series matches the statistical properties of the real time-series; 2) diversity, which quantifies the variability and richness of the synthetic time series; and 3) utility, which assesses the effectiveness of the synthetic time series in improving the prediction for positive health events with minimal records, under two evaluation scenarios (overall health events prediction based on instance and future health events prediction based on time). More details on these metrics and their implementation were provided in Figure 2C, 2D, 2E and Supplementary Methods 3.

### 2.4 Comparison with other generative models

In this study, we compared the performance of our SynLS with two state-of-the-art time series generative models. The training details and hyperparameters of these models were present in Supplementary Method 4.

1. TimeGAN ^30^ builds upon the GAN architecture, consisting of four neural networks — an embedding function, recovery function, sequence generator, and sequence discriminator. The embedding function maps raw time series into a latent space, providing inputs for the sequence generator; the sequence generator then creates new time series based on vectors from the latent space; the discriminator evaluates whether the generated time series is real or synthetic; and the recovery function maps the generated time series back to the original data space for quality evaluation. These are jointly trained to simultaneously encode features, generate representations, and iterate across time.
2. TimeVAE ^31^ presents an alternative approach for synthesizing time series from complex distributions based on VAE architecture. It comprises two primary components: the encoder and the decoder. The encoder employs convolutional and fully connected linear layers to map the input dataset to a latent space, where the latent vector follows a multivariate Gaussian distribution. The decoder takes the sampled latent vector, applies a fully connected linear layer and transposed convolutional layers to generate the synthesized output time-series. Together, these components collaborate to synthesize time-series, with the encoder extracting crucial features and the decoder reconstructing the time series based on the encoded information.

## 3. Results

### 3.1 SynLS generates high-fidelity and high-diversity time series data

To establish the reliability of our data, we first evaluated fidelity by comparing equal quantity of real and synthetic time series generated by SynLS. We analyzed the temporal trajectory (mean and standard deviation) for real and synthetic time series across four livestock wearables datasets. As illustrated in Figure 3A, SynLS is capable of synthesizing time series data highly similar to real time series data, preserving temporal dynamics patterns. For example, in dataset D1, characteristic physiological signatures near the estrus and calving periods—specifically a slight decrease in rumination time and a slight increase in activity time—were accurately captured in the synthetic time-series data. To quantitatively verify similarity, we utilized Dynamic Time Warping to measure the distance between the real and synthetic time series data (Supplemental Figure 3). The results reveal low distances across all datasets (D1 with maximum distance of 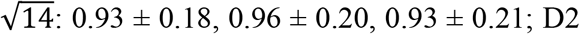 with maximum distance of 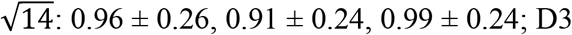 with maximum distance of 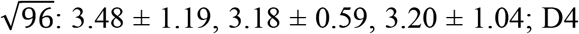 with maximum distance of 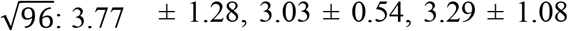; Disease, Estrus, and Calving group, respectively). These consistently low distances strongly confirm the model’s ability to generate high-fidelity livestock wearable sensors time series data.

**Figure 3.**
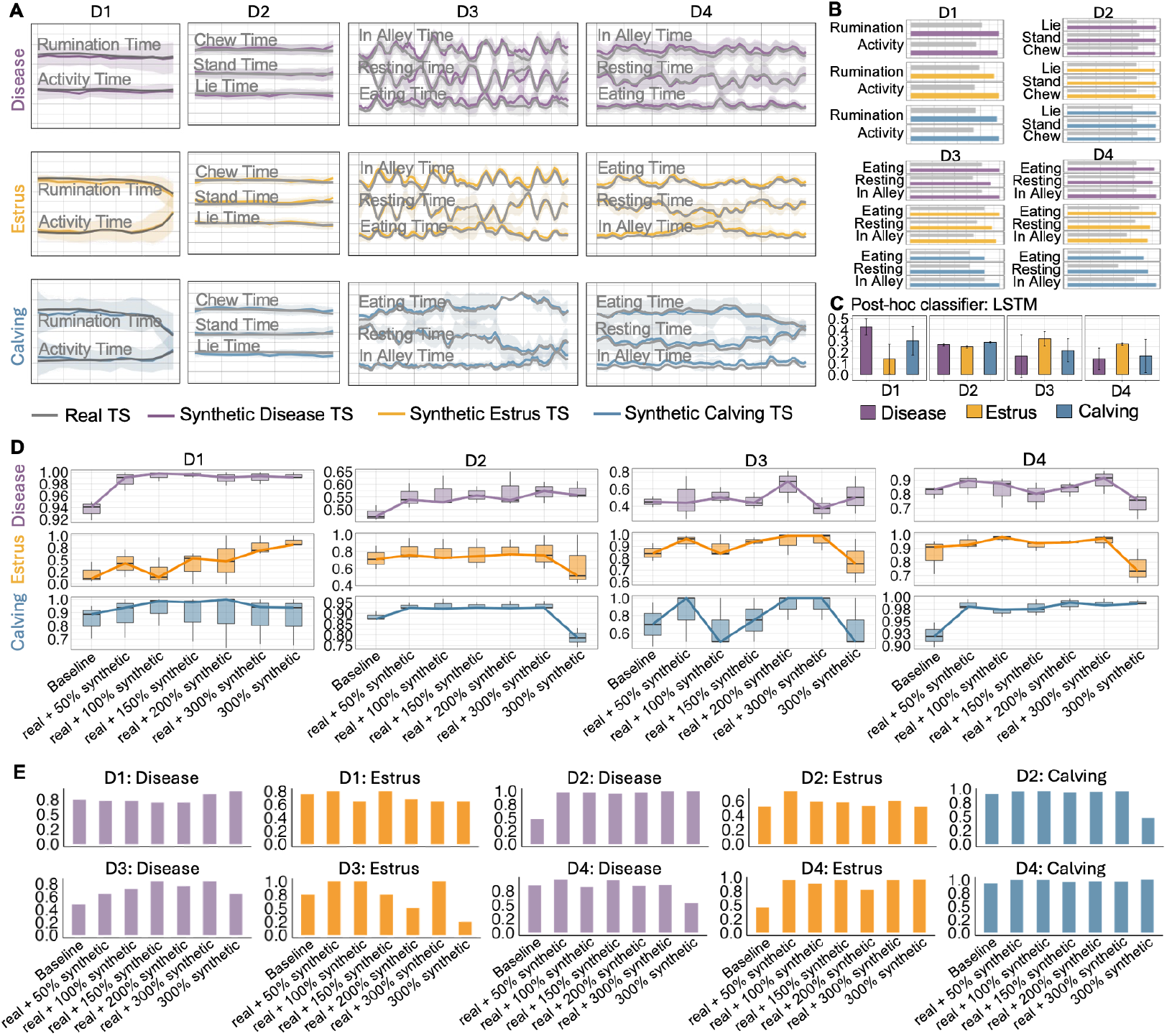
Results of SynLS’s fidelity, diversity and and utility. A. Comparison of the distribution of values at each timepoint (mean ± standard deviation) between real and synthetic wearable TS generated by SynLS. The mean value of the real/ synthetic feature at each timepoint is plotted by the solid line, with the shaded area indicating ±1 standard deviation. B. The spectral entropy describing diversity for real and synthetic wearable TS generated by SynLS. C. Discriminative score, defined as |0.5 − classifier’s accuracy|, for distinguishing real and synthetic wearable TS using Bidirectional LSTM as post-hoc classifier. D. Predictive performance (AUROCs) for each type of health event in each database with different augmentation ratios, under the overall health event prediction evaluation. E. Predictive performance (AUROCs) for each type of health event in each database with different augmentation ratios, under the future health event prediction evaluation.

Next, to assess the distribution similarity beyond time dynamics, we projected the real and synthetic time series data in a two-dimensional space using t-Distributed Stochastic Neighbor Embedding (t-SNE) algorithm (Supplemental Figure 4). We found that the distributions highly overlapped across all datasets without showing separated clusters. Interestingly, we observed that the diversity of two-dimensional spatial distribution of synthetic time series data higher than that of real time series data in most datasets. To more rigourously compare the diversity, we calculated the spectral entropy of real and synthetic time series data for each dataset and found that synthetic time series data exhibited greater diversity than real time series data, with an average increase of 29.7% across all datasets (Figure 3B; specific spectral entropy value in each database was described in Supplemental Table 4). This confirmed the model can synthesize time series data with greater diversity than the real time series data.

Finally, we evaluated the fidelity of synthetic data by employing a widely-used post-hoc discriminator, a 2-layer Bidirectional LSTM with 20 units in each layer ^31^. The discriminator’s performance was quantified using a discrimination score, which was defined as the absolute value of 0.5 minus the predictive accuracy. Results shown in Figures 3C revealed that, except for the Disease and Calving groups at D1 and Estrus group at D2 (with the average discrimination score of 0.43, 0.30 and 0.32, respectively), the average discrimination score did not exceed 0.30 in all other databases. These results demonstrates that, regardless of the varied length of the input time series, SynLS effectively generate realistic sequences that reassemble real data, making them challenging for discriminators to differentiate.

### 3.2 SynLS enhances predictive performance for livestock health events

Beyond fidelity and diversity, the ultimate utility of synthetic data is its ability to improve downstream predictive performance in precision livestock and medical research and applications. A critical real-world challenge is the severe shortage of data for positive events (e.g., disease onset, Supplemental Figure 1C), which complicates machine learning (ML) modeling and evaluation. To rigourously evaluate whether augmenting datasets with synthetic instances enhance the detection performance for the rare but critical events, we consider two evaluation scenarios: 1) overall health events prediction based on instance, 2) and future health events prediction based on timestamps (Figure 2C, 2D, Supplementary Methods 3). In each of these two evaluation scenarios, we incrementally increased the proportion of synthetic time series data (augmentation ratio) in the training dataset (0% as baseline: Train on Real, Test on Real (TRTR), 50%, 100%, 150%, 200% and 300%: Train on Synthetic and Real, Test on Real (TSRTR), solely 300% synthetic: Train on Synthetic, Test on Real (TSTR)) and tested on real time-series. A 2-layer bidirectional LSTM (with 20 units in each layer) with a single-layer fully connected head classifier ^48^ was used to predict each health event (diseases, estrus, and calving).

To ensure robust and unbiased evaluations, we implemented 3-fold cross-validation repeated three times (Figure 2C). The results (Figure 3D, specific prediction performance was described in Supplemental Table 5) indicated that augmenting training datasets with synthetic data (TSRTR) from SynLS consistently improved prediction performance compared to the baseline scenario (TRTR) across health events and datasets. Specifically, the Area Under the Receiver Operating Characteristic curve (AUROC) increased from 0.94 to 0.99, from 0.21 to 0.53, and from 0.85 to 0.88, for disease event, estrus event, and calving event, respectively in D1; from 0.48 to 0.56, from 0.72 to 0.77, and from 0.89 to 0.91, for disease event, estrus event, and calving event, respectively in D2; from 0.45 to 0.50, from 0.85 to 0.91, and from 0.70 to 0.79, for disease event, estrus event, and calving event, respectively in D3; and from 0.82 to 0.83, from 0.86 to 0.92, and from 0.92 to 0.98, for disease event, estrus event, and calving event, respectively in D4. The predictive recalls (with the cutoff point of 0.5) for each health event under 3-fold cross-validation was shown in Supplemental Figure 5. These consistent performance enhancements across multiple datasets and health events demonstrated that the synthetic data generated from SynLS effectively improve the detection of critical health events in livestock with ML models by addressing the data scarcity issue.

To evaluate the practical utility of synthetic data in real-world scenarios with limited historical data,, we partitioned each real dataset chronologically into historical (60%) and prospective (40%) segments (Figure 2D). Predictions of future events were then assessed using the previously defined evaluation frameworks baseline (TRTR), synthetic and real combined (TSRTR), and exclusively synthetic (TSTR). Following this partition, datasets D1 and D3 had no calving events in the prospective portion, so we only assessed disease and estrus events in these datasets. The results showed that the majority of models trained under TSRTR showed higher average AUROC than the baseline (disease group in D2: 0.47 vs 0.97; estrus group in D2: 0.57 vs 0.60; calving group in D2: 0.94 vs 0.95; disease group in D3: 0.48 vs 0.76; estrus group in D3: 0.75 vs 0.85; estrus group in D4: 0.45 vs 0.90; calving group in D4: 0.93 vs 0.98; baseline vs TRSTR), except for the disease group in D1 (0.78 vs 0.73; baseline vs TRSTR), estrus group in D1 (0.80 vs 0.71; baseline vs TRSTR), and disease group in D4 (0.89 vs 0.88; baseline vs TRSTR) (Figure 3E, Supplemental Table 6). The predictive recalls (with the cutoff point of 0.5) for each health event was shown in Supplemental Figure 6. Collectively, these results indicated that SynLS-generated synthetic data can address the historical data scarcity issue and enhance the model’s ability to predict future livestock health events. Consequently, SynLS holds promising for reducing the required volumn of real data while preserving adequate predictive accuracy.

A critical consideration for practical implementation of synthetic data augmentation is determining the ideal proportion of synthetic to real data that maximizes performance gains without introducing artificial biases. We demonstrated that SynLS achieved consistent and improved performance across most datasets with multiple augmentation ratios — as the ratio gradually increased from 50% to 300%, the model’s AUROC and recall either improved or remained stable under both evaluation scenarios (Figure 3D, 3E, Supplemental Figure 5, 6). However, when models were trained exclusively with 300% synthetic data (TSTR scenario), the predictive performance declined compared to the baseline in certain datasets (i.e., from 0.74 to 0.64 for the estrus group in D2; from 0.89 to 0.55 for the calving group in D2; from 0.62 to 0.54 for the disease group in D3; from 0.80 to 0.68 for the estrus group in D3; from 0.67 to 0.61 for the disease group in D4; from 0.85 to 0.79 for the estrus group in D4; AUROC in the evaluation for overall health events prediction; Supplemental Table 5 and 6). These results indicated that downstream classification tasks achieve robust and improved prediction performance when trained on a combination of real data combined with a moderate proportion of synthetic data, rather than relying exclusively on synthetic data.

### 3.3 SynLS outperforms existing generative approaches in both quality metrics and downstream task performance

To further evaluate the effectiveness of our proposed approach, we compared SynLS-generated synthetic time series with those produced by two established generative models, TimeGAN and TimeVAE. Results demonstrated that SynLS consistently generated more beneficial synthetic data for downstream predictive tasks, outperforming both TimeGAN and TimeVAE in terms of AUROC and recall across most datasets (Figures 4A, 4B). The performance differences were particularly striking for disease detection in dataset D4, where SynLS achieved an average AUC of 0.83 and recall of 0.68 for overall disease prediction, significantly outperforming TimeGAN by 35.6% and 298.4%, respectively (both *p* < 0.05; Figure 4A, Supplemental Figure 7), and TimeVAE by 39.2% and 32.8%, respectively (both *p* < 0.05; Figure 4A, Supplemental Figure 7). Similarly, SynLS dramatically surpassed both TimeGAN and TimeVAE for future disease group prediction on D2 (AUC: 0.98 vs 0.63 vs 0.44; recall: 0.97 vs 0.25 vs 0.19; SynLS vs TimeGAN vs TimeVAE; Figure 4B, Supplemental Figure 8). We additionally assessed computational efficiency by comparing the training and inference speeds among three generation models. We found that SynLS showed similar speed with TimeVAE and much higher speed than TimeGAN across the four datasets (Supplemental Figure 9). This suggested that our model, through the integration of diffusion framework and transformer mechanism, not only generates high-quality data for improved predictive utility, but also effectively alleviates the challenge of prolonged training and inference times commonly associated with traditional GANs and VAEs.

**Figure 4.**
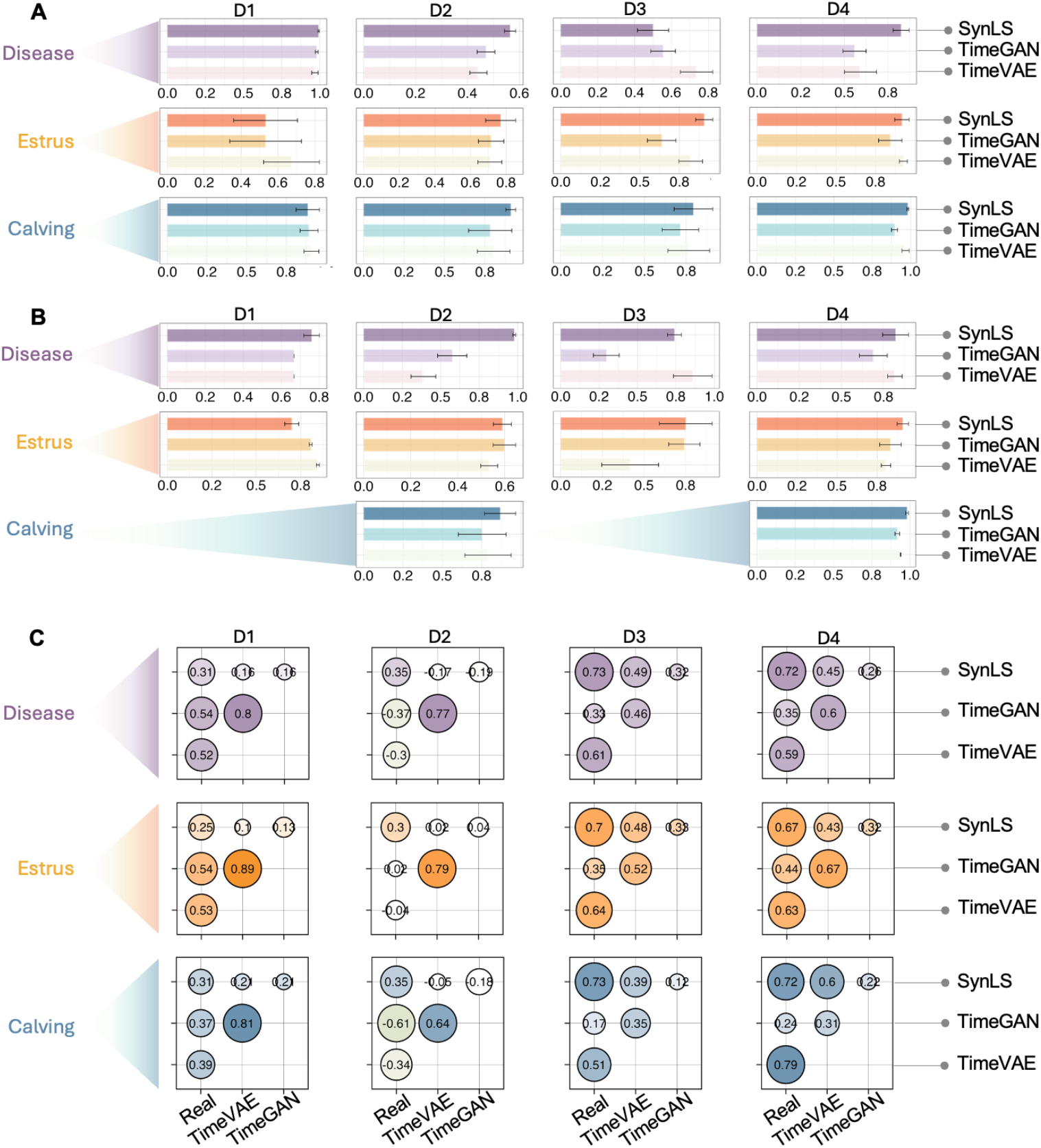
Comparison between SynLS with two alternatives. A. Average predictive performance (AUROCs) for each type of health event in each database, using synthetic data generated by SynLS, TimeGAN and TimeVAE, across different augmentation ratios ranging from 50% to 300%, under the overall health event prediction evaluation. B. Average predictive performance (AUROCs) for each type of health event in each database, using synthetic data generated by SynLS, TimeGAN and TimeVAE, across different augmentation ratios ranging from 50% to 300%, under the future health event prediction evaluation. C. Average spearman pairwise correlations between real wearable time series data and synthetic data generated by SynLS, TimeGAN and TimeVAE, in each database.

To evaluate how differently the synthetic data from TimeGAN, TimeVAE and SynLS can model the real wearable time series data, we compared Spearman correlations coefficient between synthestic and corresponding real data among three methods. We categorized the correlation coefficients into six levels: strong negative ([− 1, − 0.5)), middle negative ([− 0.5, − 0.3)), low negative ([− 0.3, − 0.1)), no correlation ([− 0.1, 0.1)), low positive ([0.1, 0.3)), middle positive ([0.3, 0.5)), and strong positive ([0.5, 1)) ^20^. As observed, SynLS significantly outperformed TimeGAN and TimeVAE in reconstructing temporal dynamics of real data (Figure 4C). For dataset D2, SynLS consistently demonstrated moderate positive correlations between its real and synthetic time series (0.3 to 0.35), in contrast to TimeGAN and TimeVAE, which displayed a range from no correlation to strong negative correlations (−0.61 to 0.02 for TimeGAN, -0.34 to -0.04 for TimeVAE). Especially for TimeGAN, it displayed significantly lower quantitative metrics than both SynLS and TimeVAE on datasets D3 and D4. This correlation analysis provide evidence that SynLS more effectively captures and reproduces underlying temporal patterns in wearable time series data than other commonly used methods.

### 3.4 Extending SynLS to raw wearables accelerometry data for improved behaviors classification

Having demonstrated effectiveness of SynLS for high-quality aggregated livestock sensor data aforementioned, we extended our investigation to evaluate its capabilities to generate high-quality raw accelerometry data and the performance of downstream task, behavior classification based on the raw accelerometry data. We collected new datasets including two livestock 3-axis accelerometry datasets and two human 3-axis accelerometry datasets (Figure 1A, 1B, Supplemental Table 2 and 3). For livestock datasets, we evaluated the model performance under 3-fold cross-validation based on instance. For human datasets, we split the datasets into one training set and one validation set as labelled by the authors ^44,45,46^.

Our findings indicated that increasing the proportion of synthetic wearables raw 3-axis accelerometry data could aid in enhancing behavior classification tasks for all the livestock and humans datasets (Figure 5). Similar to the patterns observed in health event prediction tasks, most datasets exhibited consistently enhanced (or at least stable) classification accuracies when we increased the augmentation ratios. For instance, at a 200% augmentation ratio compared to the baseline, the datasets from Ito et al. and Erik et al. showed improvements in classification accuracy: grazing accuracy increased by 22.2% and 19.4%, moving by 5.3% and decreased by 6.9%, resting by 42.7% and 71.4%, and rumination by 4.5% and 80.0%, respectively. Furthermore, when models were trained exclusively on synthetic data, their performance on real-world data might exhibit a notable decline, mirroring the challenges observed in health event prediction tasks. To illustrate, in the second human behavior dataset provided by Tonkin et al., the classification accuracy for sitting, standing, and kneeling experienced significant drops — from baseline values of 0.16 to 0.01, 0.31 to 0.14, and 0.95 to 0.51, respectively. The specific classification performance was present in Supplemental Table 7-9. These findings demonstrated that SynLS is capable to synthesize high-quality raw multi-dimensional accelerometer data with complex frequency characteristics and enhance downstream tasks, underscoring that SynLS offers a robust, versatile solution for synthetic data generation across the full spectrum of wearable sensor applications in precision monitoring, incuding human medicine.

**Figure 5.**
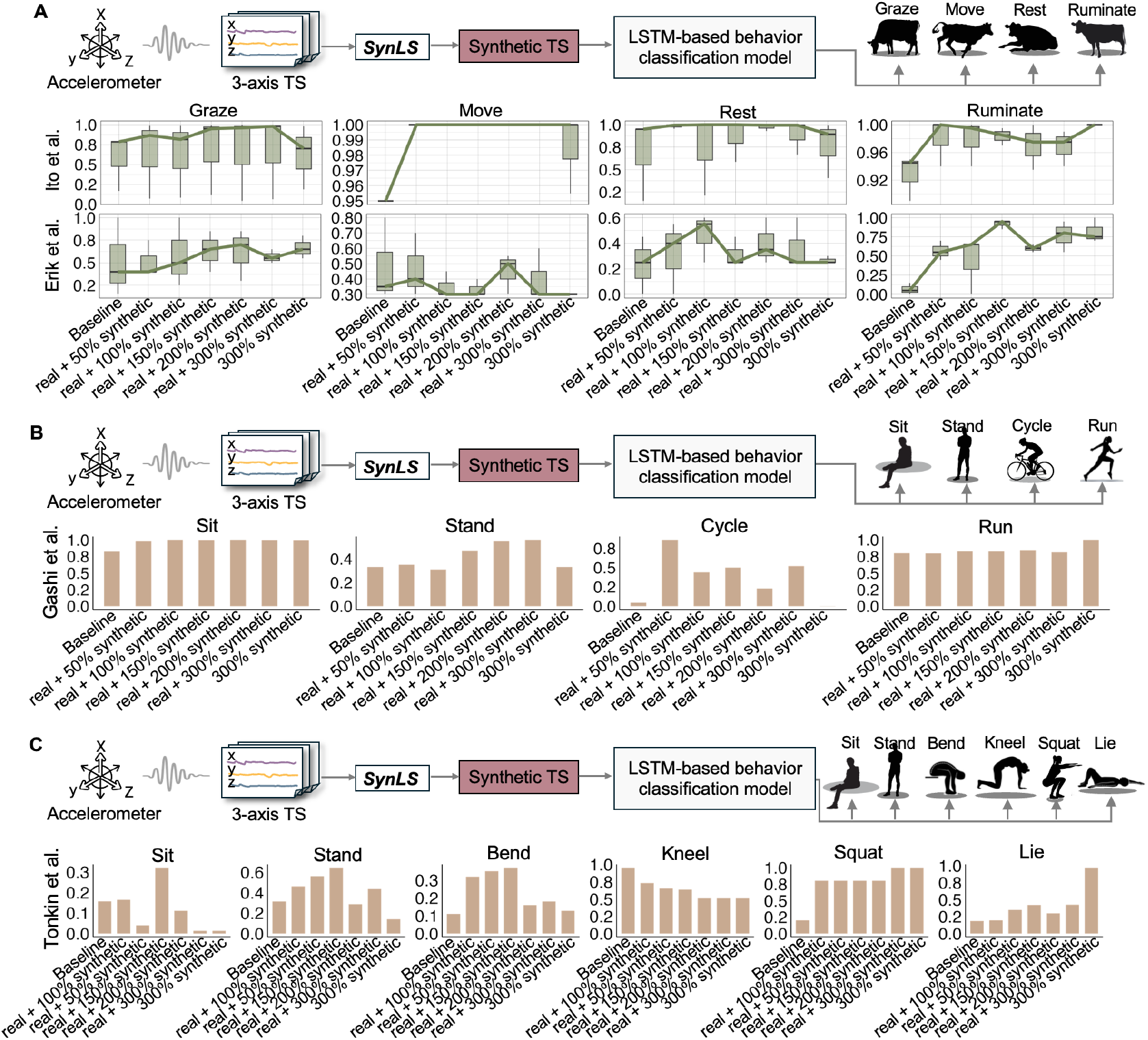
Results of SynLS’s utility in extension 3-axis accelerometry databases. A. Illustration of the livestock behavior (grazing, moving, resting and rumination) classification using synthetic data generated by SynLS and classification accuracy for each type of behavior in each livestock database with different augmentation ratios. B. Illustration of the human behavior (sitting, standing, cycling and running) classification using synthetic data generated by SynLS and classification accuracy for each type of behavior in one human database (Gashi et al.,) with different augmentation ratios. C. Illustration of the human behavior (sitting, standing, bending, kneeling, squatting and lying) classification using synthetic data generated by SynLS and classification accuracy for each type of behavior in one human database (Tonkin et al.,) with different augmentation ratios.

## 4. Discussion

In this work, we introduced SynLS, a pioneering generative deep learning model engineered to enhance health monitoring (especially for animals) by synthesizing highly realistic wearable sensor data. The innovative essence of SynLS lies in its ability to replicate intricate sensor time series data with remarkable cost-effectiveness and minimal real data requirements. This model significantly improves some downstream analyses, like health prediction (with an average increased AUC of 22.7% across datasets and augmentation ratios) and behavior classification (with an average increased AUC of 49.1% across datasets and augmentation ratios), even when only utilizing lightweight 2-layer bidirectional LSTM classifiers. SynLS provides a robust solution for improving real-time livestock health monitoring, offering implications for veterinary medicine and the broader medical sector.

The major contribution of our work is the development of a diffusion-transformer generative deep learning framework. This framework effectively addresses the challenges of wearable sensor data, including its varied length, multiple dimensions, high diversity, high noise, periodicity and trend, to generate realistic synthetic time series data. Experimental results demonstrated that downstream classifiers trained with a certain proportion of SynLS-generated synthetic data consistently outperformed both the baseline trained exclusively on real data and classifiers incorporating synthetic data from traditional GAN-based and VAE-based models (Figure 3, Figure 4). The significant improvement might be partially rooted in several advantages from DDPM framework and transformer model. The generation process of DDPM is inherently a multi-scale learning procedure ^32^, enabling the generative model to capture coarse long-term/global trends in early denoising stages and refine periodic and short-term fluctuations in later stages. Simultaneously, the nature of denoising process of DDPM may provide robustness against noise ^32^. The integration of transformer probably further enhances DDPM’s capabilities: positional encoding preserves temporal order, while the self-attention mechanism explicitly captures long-range dependencies ^49^, aiding DDPM in better modeling trends and cyclical patterns.

The second major contribution of our work lies in the demonstrated extensibility of our framework to raw triaxial accelerometer data, significantly enhancing downstream behavior classification tasks (Figure 5). Experimental results showed that our model effectively generated realistic synthetic data even from low-level sensor inputs, which are typically challenging due to their high noise levels and complex spatiotemporal patterns ^38,39,40^. This capability holds significant potential for broad medical research, where raw sensor data from wearable devices is increasingly used for continuous health monitoring and early disease detection. By generating high-fidelity synthetic raw accelerometry data, our framework shows the potential address critical challenges such as data scarcity and privacy concerns ^20,50^, enabling the development of robust diagnostic models without compromising patient confidentiality. This aligns with the growing demand for scalable and privacy-preserving solutions in medical AI research ^48,51^.

Our results revealed some interesting patterns in terms of the combination of real and synthetic time series data. In most cases, training with a combination of both outperformed baselines using only real data. In-depth analysis revealed that diffusion-generated synthetic data exhibited greater diversity (Figure 3B, Supplemental Figure 4), potential enabling downstream models to learn boarder decision boundaries and enhancing generalization capability ^52^. However, relying exclusively on synthetic data posed risks, as downstream tasks were prone to collapse (Figure 3D and 3E, Supplemental Figure 5 and 6). This highlights a key limitation — despite the effectiveness of our generative model, synthetic data cannot fully replace real data in all scenarios ^53^. Continued real data collection remains essential for robust model performance.

Our study still has several limitations. First, although our proposed model can capture variations in wearables time series, it does not explicitly condition on specific health events, which could improve its ability to generate more contextually relevant data and enhance adaptability to diverse clinical scenarios ^56^. Second, our model does not incorporate discrete animal-level variables, which are typically present alongside continuous time seires in real-world datasets and may impact health outcomes ^20^. Third, our model does not yet integrate multimodal data generation for animal health monitoring, including both time series data and images ^57,58^. Future work should incorporate discrete animal-level variables, integrate multimodal data, and condition on health events to enhance the capabilities of generative models in the veterinary medicine and even in human medicine.

In conclusion, we have developed SynLS, a time series generation model for livestock wearable sensors data based on the diffusion-transformer framework. SynLS’s architecture incorporates several innovations to overcome key challenges inherent in wearable (especially for animals) sensor data: a novel diffusion process optimized for multi-scale temporal patterns, and transformer-based encoding that preserves long-range dependencies in physiological signals. By demonstrating that carefully designed usage strategy of synthetic data can dramatically enhance classifier performance even with lightweight models, SynLS advances the feasibility of accurate, real-time health monitoring in resource-constrained livestock or human health monitoring environments. This work establishes a foundational framework for health monitoring data supplementation with immediate practical utility while opening promising research directions that could enable more efficient and effective disease detection across species.

